# A new story of four Hexapoda classes: Protura as the sister to all other hexapods

**DOI:** 10.1101/2024.01.08.574592

**Authors:** Shiyu Du, Erik Tihelka, Daoyuan Yu, Wan-Jun Chen, Yun Bu, Chenyang Cai, Michael S. Engel, Yun-Xia Luan, Feng Zhang

## Abstract

Insects represent the most diverse animal group, yet previous phylogenetic analyses based on the morphological and molecular data have failed to agree on the evolutionary relationships of early insects and their six-legged relatives (together constituting the clade Hexapoda). In particular, the phylogenetic positions of the three early-diverging hexapod groups, the coneheads (Protura), springtails (Collembola), and two-pronged bristletails (Diplura), have been debated for over a century, with alternative topologies implying drastically different scenarios of the evolution of the insect body plan and hexapod terrestrialisation. We addressed this issue by sampling of all hexapod orders, and experimented with a broad range of across-site compositional heterogeneous models designed to tackle ancient divergences. Our analyses support Protura as the earliest-diverging hexapod lineage (Protura-sister) and Collembola as a sister group to the Diplura, a clade we refer to as ‘Antennomusculata’ characterised by the shared possession of internal muscles in the antennal flagellum. The universally recognized ‘Ellipura’ hypothesis is recovered under the site-homogenous LG model. Our cross-validation analysis shows that the CAT-GTR model that recovers Protura-sister fits significantly better than homogenous model. Furthermore, as a very unusual group, Protura as the first diverging lineage of hexapods is also supported by other lines of evidence, such as mitogenomics, comparative embryology, and sperm morphology. The backbone phylogeny of hexapods recovered in this study will facilitate the exploration of the underpinnings of hexapod terrestrialisation and mega-diversity.

## INTRODUCTION

Insects represent the most prolific radiation in the animal kingdom, accounting for over half of all described metazoan species^1^. Winged insects came to dominate most terrestrial ecosystems by the late Carboniferous, over 310 Mya [million years ago]^2^. Partly due to its great antiquity, the origins of insect mega-diversity remain elusive. Current hypotheses tie the radiation of insects with their geological age, diversification rate, critical anatomical innovations, ecosystem change, and/or dietary breadth^3–7^. As the closest relatives of insects, the non-insect hexapods play a pivotal role in understanding the unparalleled evolutionary success of six-legged life^8,9^. This group comprises small-bodied, elusive terrestrial arthropods with pronounced adaptations to a soil-dwelling lifestyle. Unlike insects, these ‘basal’ hexapod clades account for <1% of animal diversity, with some 10,800 species described to date^10–12^. This group includes the comparatively species-poor and blind Protura (coneheads), the similarly speciose Diplura (two-pronged bristletails), and the considerably more diverse Collembola (springtails) armed with a characteristic abdominal jumping apparatus that gave them their name^13^. Together with insects, they constitute the clade Hexapoda^8,14^.

The availability of genome-scale datasets has helped settle numerous historical conundrums in insect phylogeny over the last two decades^8,15,16^. The dawn of the phylogenomic era has confirmed the monophyly of Hexapoda and elucidated the group’s closest relatives^8,17,18^. While traditional morphological studies considered hexapods as close relatives of myriapods^19^, molecular datasets have revealed that the group is nested within the ‘crustaceans’, as sister group to the enigmatic clade Remipedia, which inhabits flooded coastal caves^8,18,20,21^. These results backdate the origin of crown-group insects to the Silurian–Cambrian^8,22,23^ and imply that insect diversification was preceded by a terrestrialisation event^18^. However, remipedes possess numerous specializations for aquatic life and so there remains some morphological differences between them and modern hexapods. The early evolution of the hexapods thus remains veiled in mystery, not only because of the extreme scarcity of hexapod fossils before the Late Carboniferous^24^, but also because the relationships among the earliest-diverging hexapods have proven resistant to resolution, whether interrogated with morphological^14,25,26-28^, single-gene^29,30^, mitochondrial^31,32^, phylogenomic data^8,17,18,33^, as well as combined analyses^34-36^. Recent studies are mostly split between favouring a clade of Protura + Collembola (the ‘Ellipura’ hypothesis)^8^, Protura + Diplura (the ‘Nonoculata’ hypothesis)^17,27,37–40^, or Diplura + Collembola^20,41^, and Diplura + Insecta (the ‘Cercophora’ hypothesis)^42,43^. Earlier morphological studies have cautiously treated the ‘basal’ hexapod clades as a single clade, ‘Entognatha’^14,44^, while others maintain that the ‘basal’ hexapods form a paraphyletic grade^45^. Traditional morphological studies, conducted since the 19^th^ century^46,47^, are confounded by the ‘basal’ hexapod’s extreme specialisations for life in the soil, which makes inferring homologous characters challenging^48,49^. Molecular studies are complicated by the rarity and small size of many morphologically peculiar ‘basal’ hexapod groups, which have so far been sampled only sparsely in phylogenomic studies. Moreover, the great antiquity of the divergence between the ‘basal’ hexapods and crown-group insects represents a formidable challenge to conventional molecular phylogenetic methods, as ancient rapid divergences often induce phylogenetic artifacts such as long-branch attraction^50,51^.

Here we address the problem of insect origins by increasing the taxon and gene sampling of overlooked groups. We sequenced the transcriptome for a second proturan species, belonging to the genus *Sinentomon*^29–31^, along with two new transcriptomes for dipluran species. We employ a variety of analytical approaches to account for common sources of error in phylogenomics, interrogate the robustness of the results, and interpret them with respect to the origin of insect body plan and hexapod terrestrialisation.

## RESULTS

### Genomic sequencing and matrix assembly

We sequenced the transcriptome of the proturan *Sinentomon erythranum* (SRX480876; Fig. 1). *S. erythranum* is a member of the rare monogeneric family Sinentomidae endemic to eastern Asia. This group was not discovered until the 1960s^52^ and its phylogenetic position has stirred much controversy given the proturan’s unusual head morphology and sperm ultrastructure^53–55^. An analysis of two ribosomal RNA gene sequences recovered Sinentomidae as the earliest-diverging proturan lineage^29^, albeit substantial incongruence persists among studies^30,31,37,56,57^. We furthermore additionally sequenced two transcriptomes belonging to the diplurans *Octostigma sinensis* (SRX3641158) and *Lepidocampa weberi* (SRX3641157), representing the superfamilies Projapygoidea and Campodeoidea, respectively. Projapygoids are a presumed evolutionary link between Campodeoidea and Japygoidea^58^, but they are very rare and hard to collect for comparative studies. The interrelationships of three superfamilies and the monophyly of Diplura have been much debated. It has been suggested that diplurans may together represent a polyphyletic grade rather than a clade based on ovarian and spermatozoal characters^59,60^ albeit comparative embryological evidence and molecular evidence so far overwhelmingly supports dipluran monophyly^8,29,32,61^.

**Fig. 1.**
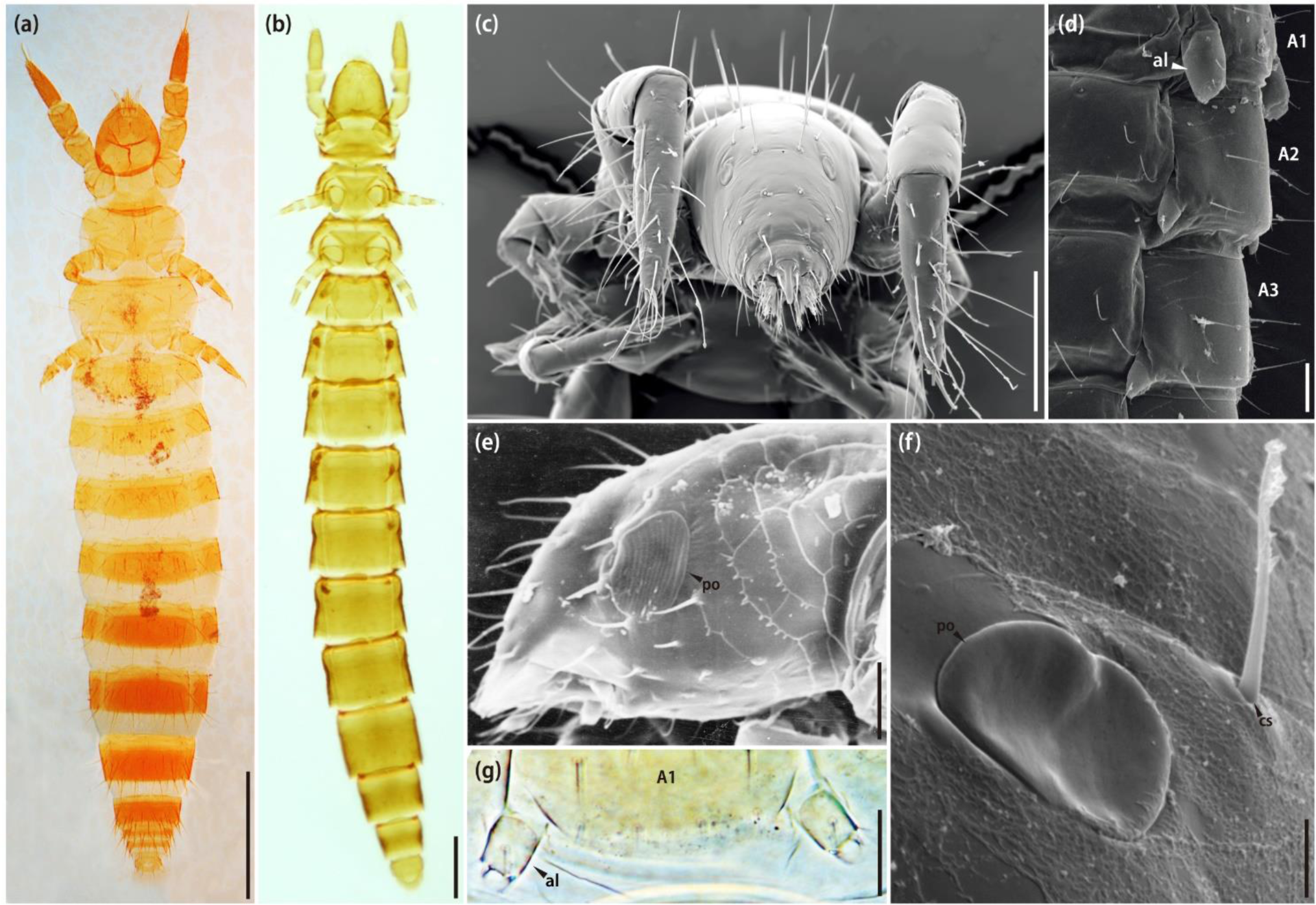
Morphology of the proturans *Acerentomon microrhinus* (Acerentomidae) and *Sinentomon erythranum* (Sinentomidae). (A) Habitus view of *A. microrhinus* under reflected light. (B) Habitus view of *S. erythranum* under reflected light. (C) Scanning electron micrograph of *A. microrhinus* head and forelegs. (D) Scanning electron micrograph of the abdomen of *A. microrhinus* in lateral view. (E) Scanning electron micrograph of *S. erythranum* head in lateral view. (F) Detail of the pseudoculus of *A. microrhinus.* (G) Abdominal legs of *S. erythranum*. Abbreviations: A1–3: abdominal segments 1; al, abdominal legs; cs, cephalic seta; po, pseudoculus. Scale: 5 μm (G); 10 μm (F); 20 μm (C, D, E); 50 μm (A, B).

We compiled genomic and transcriptomic data for 42 other hexapod species (downloaded from the NCBI; see part of METHOD DETAILS) with high near-universal single-copy orthologs gene completeness (BUSCO) scores plus three aquatic ‘crustacean’ clades (outgroups) recovered as close relatives of hexapods^18,20,21^. The inclusion of early-diverging dipluran and proturan groups is of particular relevance, as previous studies have indicated that the hexapod tree is prone to long-branch artifacts^16,20,32^, which are exacerbated by limited taxon sampling^62^. To ensure the quality of the genome/transcriptome, all species’ BUSCO assessments in this study were all above 70% (Supplementary Table 1).

Our dataset comprised a total of 48 species (including the additional three outgroups). Phylogenetic analyses were based on four amino acid (AA) alignments to explore alternative sources of phylogenomic signal. Matrix1 was generated by selecting universal single-copy orthologues (USCOs) for the 48 taxa. Trimming reduced the original dataset by 59.8% (from 1,281,520 AA sites to 515,770), and increased data occupancy from 32.68% to 66.81%. Filtering by the number of parsimony-informative sites, relative composition variability (RCV), and stationary, reversible and homogeneous (SRH) assumptions reduced the dataset by 1.7% (from 515,770 AA sites to 506,831), 20.0% (from 506,831 AA sites to 405,537), and 8.3% (from 405,537 AA sites to 371,709), respectively. TreeShrink was further used to generate a matrix with 75% completeness (the BUSCO ids and names of the putatively spurious sequence after spurious homolog identification by using TreeShrink are listed in Supplementary Table 2). In its final form, Matrix1 contained 780 loci (342,252 AA sites). Matrix2 (USCO75_abs70) was generated using genes from Matrix1 with average bootstraps support (ABS) values over 70 and consisted of 505 genes (255,095 AA sites). Subsequently, in order to detect conflicts between concatenation and coalescent-based phylogenies, Matrix1-con and Matrix2-con were generated by selecting inconsistent genes (i.e., those with gene-wise phylogenetic signal (ΔGLS)>0, or gene-wise quartet scores (ΔGQS)<0; see part of METHOD DETAILS) from Matrix1 and Matrix2, respectively. Matrix1-con consisted of 468 genes (201,896 AA sites) and Matrix2-con of 298 genes (149,903 AA sites; Supplementary Table 3).

The length, number of parsimony-informative sites, RCV values, and SRH values for every locus from each matrix were compared with a paired *t*-test (Supplementary Fig. 1). The analysis shows significant differences between Matrix1 and Matrix2, and Matrix1-con and Matrix2-con in terms of their length (*p*-value < 0.001 between Matrix1 and Matrix2, *p*-value < 0.001 between Matrix1-con and Matrix2-con) and the number of parsimony-informative sites (*p*-value < 0.001 between Matrix 1 and Matrix 2, *p*-value < 0.001 between Matrix1-con and Matrix2-con; Supplementary Fig. 1a, b). The RCV and SRH values showed no significant difference between the matrices (Supplementary Fig. 1c, d).

### Hexapod phylogeny

All our phylogenomic analyses recovered strong support for the monophyly of Collembola, Protura, Diplura, and Insecta, respectively (Bayesian Posterior Probabilities (BPP) = 1, SH-aLRT/UFBoot2 = 100/100, and ASTRAL bootstraps= 1; Fig. 2). A total of 28 ML trees and one BI tree were inferred from the four matrices (Supplementary Table 4; Supplementary Data 2) to test the effect of the substitution model on the recovered topology. Trees based on different matrices and inference models were congruent at most nodes (Fig. 2) but resulted in four different topological hypotheses (H1–4) about the relationships of the early-diverging hexapod clades (Fig. 3). Hypothesis 1 supported the placement of Collembola as sister group to the remaining hexapods (H1: ‘Collembola-first’, i.e., Collembola + (Protura + (Diplura + Insecta))). Under the second hypothesis (H2: (Collembola + Protura) + (Diplura + Insecta)), Collembola and Protura formed a monophyletic group as sister Diplura + Insecta, corresponding to the ‘Ellipura’ hypothesis^8^. Protura was inferred as the sister group to the remaining three hexapod groups in the third hypothesis (H3: ‘Protura-first’, i.e, Protura + ((Collembola + Diplura) + Insecta)). Under the fourth hypothesis (H4: (Protura + (Collembola + Diplura)) + Insecta), the clade (Protura + (Collembola + Diplura)) formed a sister group to Insecta, corresponding to the traditional concept of ‘Entognatha’^25^.

**Fig. 2.**
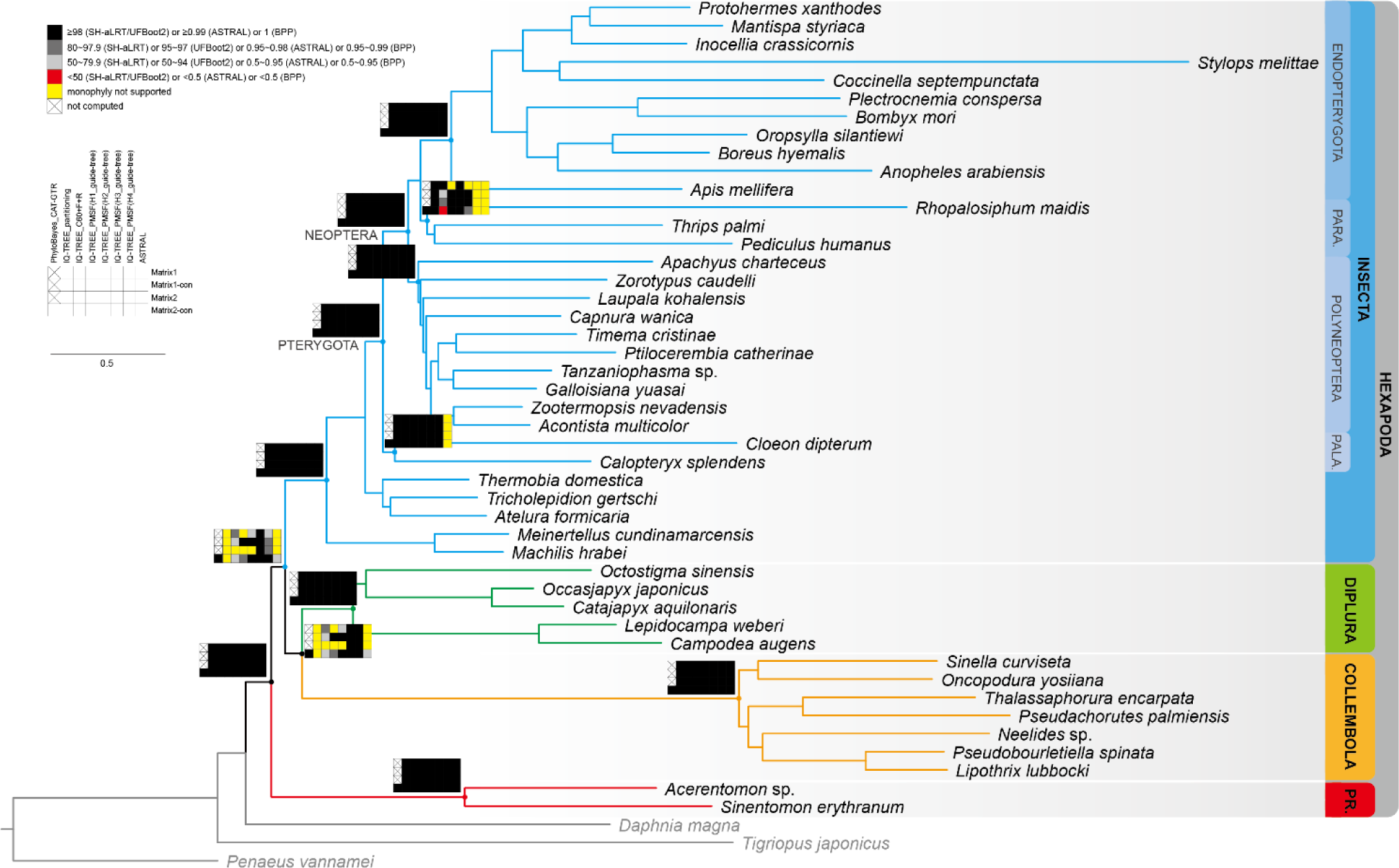
Phylogeny of the ‘basal’ hexapods. Main topology inferred from Matrix2-con using the Bayesian across-site compositional heterogeneity model CAT-GTR model implemented in PhyloBayes. Node supports from all analyses are indicated by the coloured squares (The node supports of each phylogenetic tree is shown in Supplementary Appendix A). Only the lowest support values are shown when different matrices or different models produced conflict results. Abbreviations: PARA., Paraneoptera; PALA., Palaeoptera; PR., Protura. (H1_guide-tree: Collembola + (Protura + (Diplura + Insecta)); H2_guide-tree: (Collembola + Protura) + (Diplura + Insecta); H3_guide-tree: Protura + ((Collembola + Diplura) + Insecta); H4_guide-tree: (Protura + (Collembola + Diplura)) + Insecta).

**Fig. 3.**
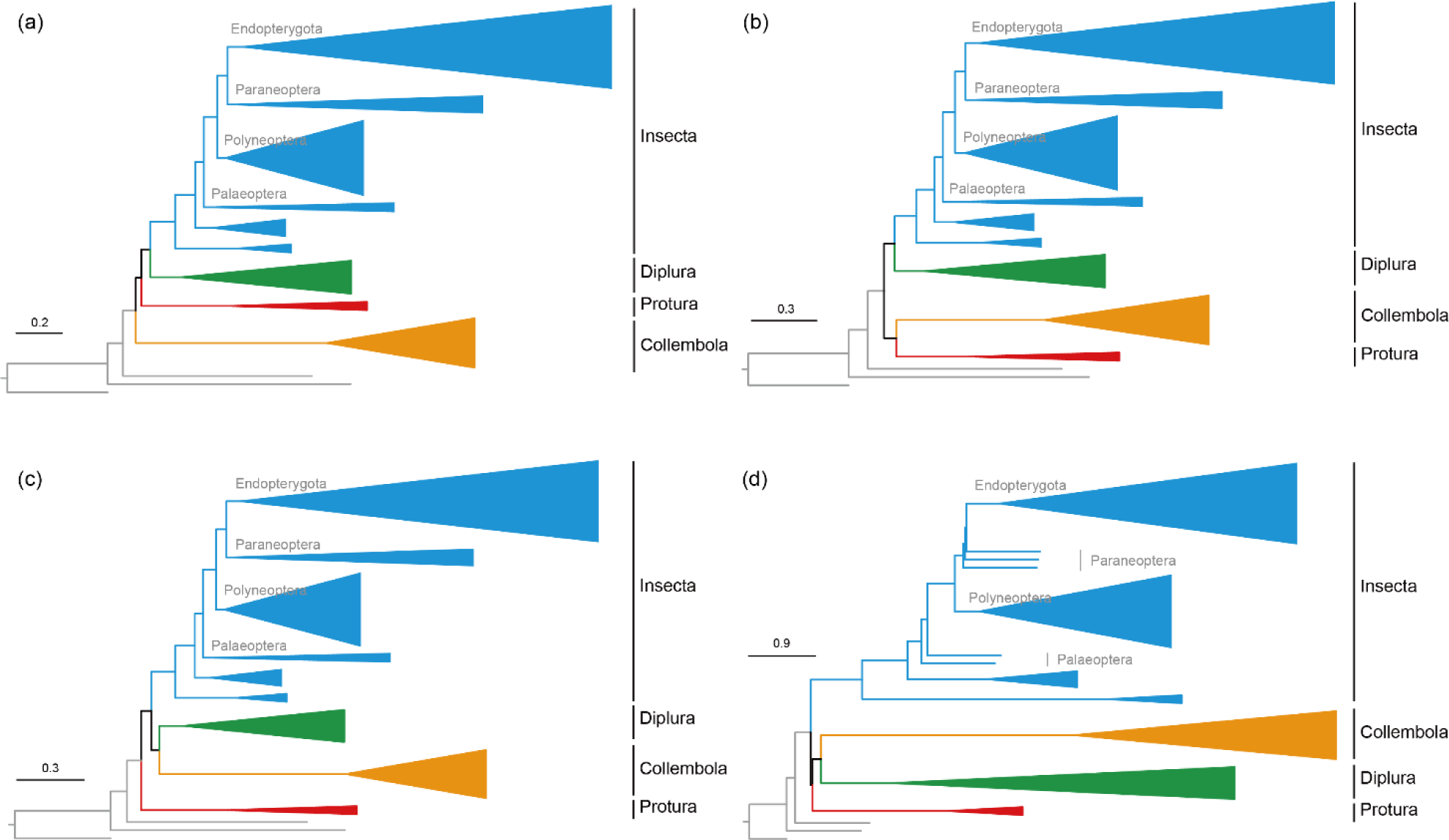
Four different topological hypotheses analysed in this study. (A) Hypotheses H1 inferred from Matrix2 using C60+F+R model implemented in IQ-TREE: Collembola + [Protura + [Diplura + Insecta]] (Collembola-first). (B) Hypotheses H2 inferred from Matrix1 using partitioned maximum likelihood model implemented in IQ-TREE: [Collembola + Protura] + [Diplura + Insecta] (the ‘Ellipura’ hypothesis). (C) Hypotheses H3 inferred from Matrix2-con using C60+F+R model implemented in IQ-TREE: Protura + [[Collembola + Diplura] + Insecta] (Protura-first). (D) Hypotheses H4 inferred from Matrix1-con using MSC model implemented in ASTRAL: [Protura + [Collembola + Diplura]] + Insecta (the ‘Entognatha’ hypothesis).

The most complex models, the finite mixture site-heterogeneous models C60+F+R and LG+PMSF(C60) and the infinite site-heterogeneous model CAT-GTR, supported H3 when Matrix1-con and Matrix2-con were analysed. Under this topology, Protura was the sister group to Diplura + Collembola and the remaining hexapods (H3). In a cross-validation test conducted on Matrix2-con, the infinite mixture model CAT-GTR fitted the dataset better than LG (cross-validation log-likelihood scores = -48079.05 ± 917.74 > -49316.14 ± 958.82; Supplementary Table 5). The Wilcoxon test analysis shows that there is a significant difference between these two models (*p*-value = 0.01469; Supplementary Fig. 2). Support for H3 declined with other finite mixture models C60+F+R and LG+PMSF(C60) that supported a broader range of topologies, with Matrix1 favouring H1 and H3, and Matrix2 H1, H2, and H3 (Supplementary Data 2). All partitioned analysis reconstructions supported topology H2 (Supplementary Table 4), while multispecies coalescent analyses of the four matrices recovered three hypotheses (H1, H3, and H4), albeit some nodes were poorly supported (Supplementary Data 2). In addition, the gene concordance factors (gCF) and the site concordance factors (sCF) were used to gain a deeper understanding of how well different genes and sites support the different hypotheses (Supplementary Data 3). For most branches in all four topologies, the gCF values are lower than the sCF values, suggesting that the sites that support these topologies are scattered across the different genes.

To test the effect of the outgroup sampling on the ingroup topology, a rooted tree without the outgroups was inferred using reversible models. Relative positions between or within the four classes in the unrooted topology are shown in Supplementary Data 4. A rooted tree (Supplementary Data 4) inferred with a non-reversible models placed ((Collembola + Protura) + Diplura) at the root, with a bootstrap value of 84. Bootstrap support for each branch is defined as the proportion of rooted bootstrap trees that have the root on that branch. Two nodes presented the bootstrap support values (Supplementary Data 4, rootstrap.nex): 69.3 for the root ((Collembola + Protura) + Diplura), and 15.3 for Collembola. These two largest bootstrap values supported the topologies H4 (‘Entognatha’) and H1 (‘Collembola-first’). Furthermore, topology tests (Supplementary Data 4, root_test.csv) provided AU *p*-values greater than 0.05 for three branches indicating them as the possible root (H4, H2, H1). Overall, all these two analyses indicate that the outgroup choice has little effect on the reconstructions of ingroup relationships.

### Evaluating alternative hypotheses and phylogenetic support

Topology tests conducted on all four matrices with the PMSF(C60) model (H3_guide-trees) and C60+F+R model using approximately the unbiased (AU), weighted Kishino-Hasegawa (WKH), and weighted Shimodaira-Hasegawa (WSH) tests. Under the PMSF(C60) model rejected hypotheses H1, H2 and H4 with strong confidence (*p* < 0.05 in most of cases) and supported hypothesis H3, with Protura as sister group to the remaining hexapods (Supplementary Table 6). But under the C60+F+R model, four matrices rejected hypothesis H4 with strong confidence (*p* < 0.05 in all cases), but only Matrix2-con supported hypothesis H3. Matrix1 and Matrix2 supported hypothesis H2 with no significant, and Matrix1-con supported hypothesis H2 with no significant (Supplementary Table 6).

To further explore the phylogenetic signal of different models and assess their impact on tree inferences considering distinct gene properties, we quantified the phylogenetic signal, or comparison of topological differences. Detailed information regarding the methods and results can be found in Supplementary Data 1.

## DISCUSSION

### Molecular and morphological congruence

As with many other ancient radiations^16^, molecular phylogenetic studies have found it challenging to elucidate the relationships of the ‘basal’ hexapod clades, which may have diverged as early as the Cambrian – Silurian^8,63^. Expanding the taxon sampling of ‘basal’ hexapods, including sequencing the transcriptome of the enigmatic *Sinentomon*, enabled us to explore various sources of phylogenomic signal and mitigate common artifacts at the base of the hexapod tree of life, which has been plagued by topological uncertainty^9,16^. We recovered four alternative topologies, corresponding to long-standing competing hypotheses regarding insect origins^8,39,64^ (Table S3; Fig. 3). Under the partitioned LG model, which supports the ‘Ellipura’ hypothesis as in Misof et al.^8^, the multispecies coalescent analyses resulted in the recovery of three hypotheses (Table S3). Moreover, the results suggest that the finite mixture site-heterogeneous models C60+F+R and LG+PMSF(C60), as well as the infinite site-heterogeneous model CAT-GTR analyses, specifically provide support for Protura as the first diverging lineage of hexapods. The question, then, is not why similar analyses give different results, but how we should interpret variation in results obtained from different analyses. The first important insights pertain to model fit. In PhyloBayes, cross-validation is a reliable and recommended approach for assessing the fit of models and is often employed to test if different substitution models significantly improve the fit to the datasets. We used cross-validation in PhyloBayes to evaluate CAT-GTR and LG models for the Matrix2-con. Our analysis revealed that CAT-GTR provided a better fit to the dataset compared to LG (Supplementary Fig. 2). The Wilcoxon test analysis indicated a significant difference between these two models. Therefore, cross-validation supports the hypothesis that the hetergenous model CAT-GTR are a better fit than the homogenous models with LG. Other topologies were supported by less well-fitting models, and by partitioned analyses, the latter of which has been shown to fit empirical data significantly less than approaches that consider heterogeneity at the site level, in most cases^65^. The second insight pertains to topology tests. We compared the four topologies on all matrices under LG+PMSF(C60) (H3 as the guide tree) and C60+F+R models using the AU, WKH, and WSH tests. All results rejected hypothesis H4 with strong confidence. Most of the results supported hypothesis H3 with strong confidence. These analyses suggest that we could recover Protura-sister over the much broader substitution model and topology test.

Proturans have long been considered as the most morphologically divergent hexapods, leading some early authors argue that they may not be related to hexapods at all^66^. The status of proturans as the earliest-diverging hexapods is further supported by a suite of morphological characters shared with myriapods and crustaceans. In proturans, the first three abdominal segments retain segmented or unsegment vestigial appendages (Fig. 1d &1g: al)^67^, a plesiomorphy shared with most myriapods and crustaceans where all trunk segments are equipped with a pair of segmented limbs^68^. These abdominal appendages have been reduced to unsegmented stubs or have been lost altogether in most hexapods^69^. A further plesiomorphic character proturans share with myriapods and crustaceans^70^, but not other hexapods, is their anamorphic postembryonic development (anamorphic development may be plesiomorphic), but it is highly variable in groups like myriapods, where epimorphic development is common (e.g., Scolopendromorpha, Geophilomorpha). That is, proturans emerge from the egg with nine abdominal segments, add a segment with the first molt and two more segments with the second molt, which results in 12 segments in the adult abdomen, including a distinct telson segment. The proturan embryonic membrane possess the ability to differentiate into the dorsal body wall, a feature shared with aquatic ‘crustaceans’ and myriapods, but not with other hexapods^71^. A further potential plesiomorphy of proturans may be the single claw (pretarsus) on each leg, while other hexapods have a pair of tarsal claws^72^. Proturans have no antennae, and they walk on four legs with the front two re-purposed as antennae, which diverges strongly from other hexapods^73^. They have no eyes, just pseudoculi, whose homology remains uncertain, which probably only sense light without forming images (Fig. 1e & 1f: po)^74^. Flagellate spermatozoa in proturans show a variable axonemal pattern, but a common, distinctive feature is the absence of central microtubules^75^. Proturans moreover possess a simplified or absent tracheal system unlike any other hexapods^76^; when tracheae are present at all, they are present as only two pairs of spiracles on the thorax^77^.

Characters supporting a Collembola + Diplura clade are fewer but include a similar process of blastokinesis^78^, and each antennal division with intrinsic musculature, whereas in the Insecta only the antennal scape possesses intrinsic muscles^26^. A close relationship between the two groups is moreover supported by some analyses of mitochondrial protein-coding genes^61^ and genomic datasets under heterogeneous models^20^. We herein propose the name ‘Antennomusculata’ for the Collembola + Diplura clade, in reference to the group’s shared antennal flagellum with intrinsic muscles.

### Implications for hexapod terrestrialisation

The terrestrialisation of hexapods, the most cryptic episode of the clade’s evolutionary history, has long remained shrouded in mystery, but equally attracted interest due to its importance for delimitating the groundplan of the ancestral hexapod. The resolution of proturans as the earliest-diverging hexapods enables to trace the sequence of character evolution and establishing homologies. Our results suggest that the last common ancestor of the hexapods was terrestrial, in contrary to some earlier hypotheses that suggested possible aquatic or semi-aquatic modes of life in early hexapod^48,79^. Fossil mycorrhizal fungi are known from the Early Devonian^80^, and molecular clock studies suggest they were present as early as the Cambrian^81^, highlighting a possible food source for early hexapods that may have facilitated their invasion of land.

A lasting contention in understanding hexapod terrestrialisation is whether adaptations for life on land were acquired in a step-wise fashion, or if the last common ancestor of Hexapoda already possessed a complex respiratory, reproductive, and sensory systems^82,83^. Some molecular and morphological studies over the past decade have argued that given their unusual organ systems, some proturan characters of the reproductive and respiratory systems may not be homologous with other hexapods and instead represent an independent ancient lineage^41,82,84^. We refrain from a more extensive discussion of ancestral hexapod traits, since some character systems are scarcely known in the ‘basal’ lineages such as Protura and Diplura. Resolution of the relationships among ‘basal’ hexapods will further facilitate ground plan comparisons with other arthropod lineages and the reinterpretation of controversial fossils^85^ that may help trace the transition of marine pancrustaceans to the terrestrial realm.

## SUPPLEMENTARY MATERIAL

Data available from the GitHub: https://github.com/xtmtd/Phylogenomics/tree/main/basal_hexapods/Supplementary_Material

## ACKNOWLEDGEMENTS

We thank two anonymous reviewers for their valuable comments. This research was funded by the National Natural Science Foundation of China (32270470, 31970434, 32170425, 31970438), and the National Science and Technology Fundamental Resources Investigation Program of China (2018FY100300).

## AUTHOR CONTRIBUTIONS

Conceptualization, S.D., Y.X.L. and F.Z.; formal analysis, S.D., F.Z. and E.T.; investigation, S.D., F.Z., CC. and E.T.; methodology, S.D., F.Z. and E.T.; transcriptome sequencing: W.J.C., Y.B. and Y.X.L.; data acquisition, S.D. and F.Z.; writing – review & editing, all authors.

## DECLARATION OF INTERESTS

The authors declare no competing interests.

## STAR METHODS

### KEY RESOURECES TABLE

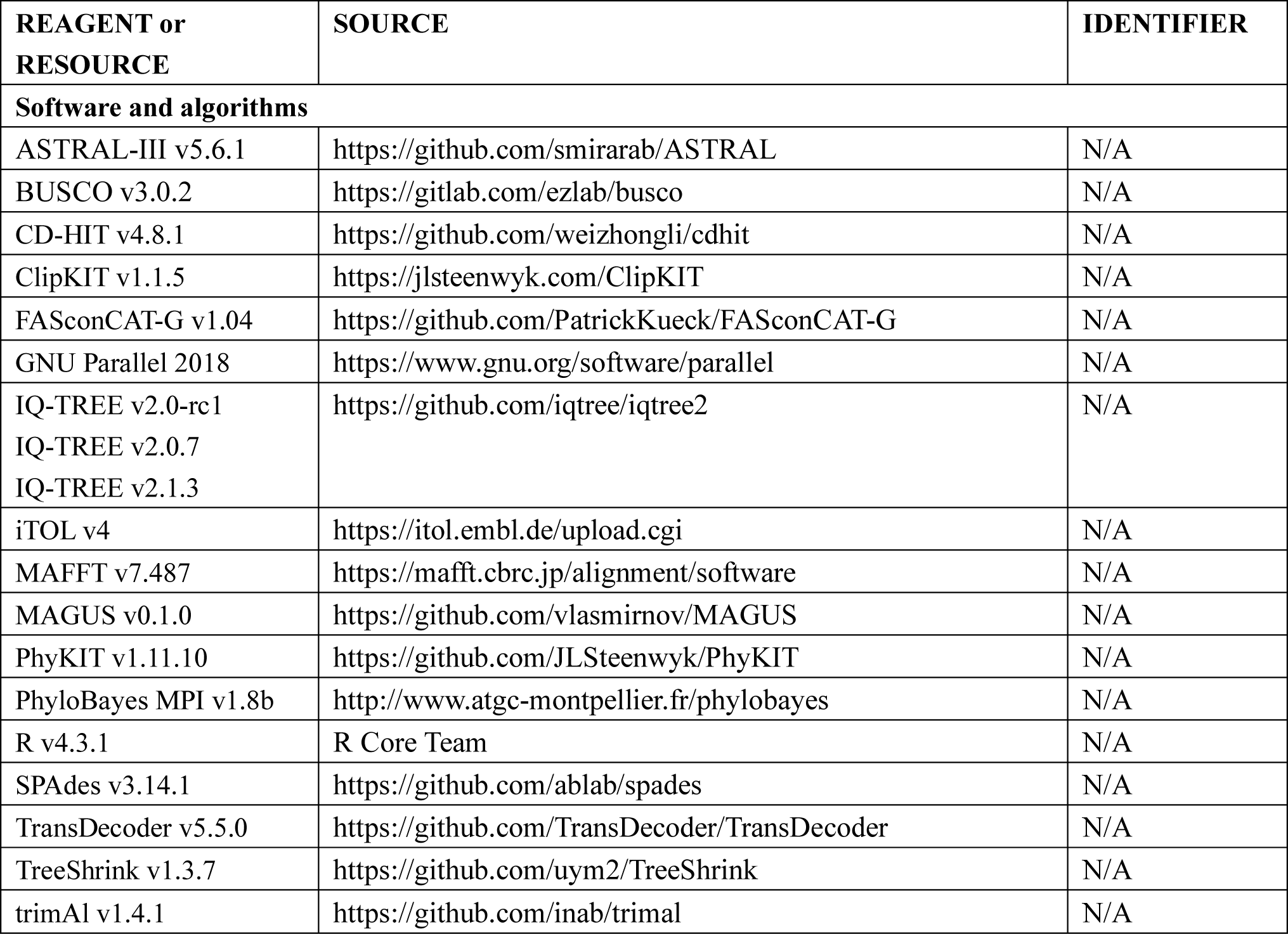

KEY RESOURCES TABLE
| REAGENT or RESOURCE | SOURCE | IDENTIFIER |
| --- | --- | --- |
| **Software and algorithms** |  |  |
| ASTRAL-III v5.6.1 | https://github.com/smirarab/ASTRAL | N/A |
| BUSCO v3.0.2 | https://gitlab.com/ezlab/busco | N/A |
| CD-HIT v4.8.1 | https://github.com/weizhongli/cdhit | N/A |
| ClipKIT v1.1.5 | https://jlsteenwyk.com/ClipKIT | N/A |
| FASconCAT-G v1.04 | https://github.com/PatrickKueck/FASconCAT-G | N/A |
| GNU Parallel 2018 | https://www.gnu.org/software/parallel | N/A |
| IQ-TREE v2.0-rc1<br>IQ-TREE v2.0.7<br>IQ-TREE v2.1.3 | https://github.com/iqtree/iqtree2 | N/A |
| iTOL v4 | https://itol.embl.de/upload.cgi | N/A |
| MAFFT v7.487 | https://mafft.cbrc.jp/alignment/software | N/A |
| MAGUS v0.1.0 | https://github.com/vlasmirnov/MAGUS | N/A |
| PhyKIT v1.11.10 | https://github.com/JLSteenwyk/PhyKIT | N/A |
| PhyloBayes MPI v1.8b | http://www.atgc-montpellier.fr/phylobayes | N/A |
| R v4.3.1 | R Core Team | N/A |
| SPAdes v3.14.1 | https://github.com/ablab/spades | N/A |
| TransDecoder v5.5.0 | https://github.com/TransDecoder/TransDecoder | N/A |
| TreeShrink v1.3.7 | https://github.com/uym2/TreeShrink | N/A |
| trimAl v1.4.1 | https://github.com/inab/trimal | N/A |

### RESOURCE AVAILABILITY

#### Lead contact

Further information and requests for resources should be directed to and will be fulfilled by the lead contact, Feng Zhang.

#### Data and code availability

All custom scripts are based on Du et al.^86^ and can be found at GitHub (https://github.com/xtmtd/Phylogenomics/tree/main/scripts and https://github.com/xtmtd/Phylogenomics/tree/main/basal_hexapods/scripts). All datasets are available at https://doi.org/XXX and are publicly available as of the date of publication. NCBI accession numbers are provided in Supplementary Table 1. Any additional information required to reanalyse the data reported in this paper is available from the lead contact upon request.

### METHOD DETAILS

#### Specimens, transcriptome sequencing, and taxon sampling

We newly sequenced three ‘basal’ hexapod transcriptomes, representing rare dipluran and proturan groups that have not been sampled previously. All three ‘basal’ hexapods were collected by YXL’s group. Specimens of *Sinentomon erythranum* Yin, 1965 were extracted from soil samples from the Tianping Mountain (Jiangsu Province, China) using Tullgren funnels. More than 200 individuals were pooled together to extract total RNA for transcriptome sequencing. The projapygoids *Octostigma sinensis* Xie & Yang, 1991 were sampled from the type locality in Zhanjiang (Guandong Province, China), a mixture of about 30 individuals was used for RNA extraction. The campodeids *Lepidocampa weberi* Oudemans, 1890 were sampled from the Shanghai Botanic Garden and the total RNA was extracted using Qiagen RNeasy Micro Kit following the manufacturer’s recommendations. Transcriptome sequencing was performed by commercial services from Beijing Genomics Institute (BGI) in Shenzhen, China using an Illumina Hiseq 2000/2500 sequencer (PE150). Raw sequencing data, and assembly accessions are provided in Supplementary Table 1.

A total of 45 hexapod species were sampled, including seven species of Collembola (representing all five orders), five diplurans (representing all three superfamilies), two proturans (representing two of three orders), and 31 insects (representing all 27 orders) (Supplementary Table 1). Care was taken to sample the three ‘basal’ hexapod groups as densely as possible. Within Insecta, only one species of each order (except for two species from Archaeognatha, three species from Zygentoma, and two species from Mecoptera) was sampled (Supplementary Table 1). Since the aim of this study is not to clarify the relationship within Insecta, the sampling of these taxa will not affect our main results and conclusion. The monophyly of hexapods has been well established^8,17,18^, three crustacean taxa were used as outgroups, following recent phylogenomic studies (e.g., Misof et al.^8^). Altogether, 48 taxa were sampled including 25 genomes and 23 transcriptomes. Publicly available genome and transcriptome assembles for 42 species were downloaded from NCBI (Supplementary Table 1). Outgroup taxa included three non-hexapod pancrustaceans based on previous phylogenomic analyses. Species names, taxonomic ranks, raw sequencing data, and assembly accessions are provided in Supplementary Table 1.

#### Genome assembly and BUSCO assessment

All paired-end reads from the three newly sequenced transcriptomes were assembled using SPAdes v3.15.5^87^. BUSCO assessments of all 48 species were conducted using the OrthoDB version 10 of the Arthropoda database (*n*=1,013) from BUSCO v3.0.2 (Supplementary Fig. 3; ^88^), with the command of ‘-m geno’. Modified the standard deviations (*σ*) of the mean USCO length to 2*σ* to be identified as ‘complete’. The BUSCO completeness values (complete and single-copy BUSCOs + complete and duplicated BUSCOs) ranged from 74.8% to 99.7% (936 ± 67.9; Supplementary Table 1).

#### Gene properties and matrix generation

Universal single-copy orthologues (USCOs) of each species were extracted, and the USCO amino acid (AA) sequences were used for subsequent analyses. Each USCO AA sequence was separately aligned using MAGUS (similar to MAFFT-linsi; ^89^). All alignments were trimmed with ClipKIT^90^ (https://jlsteenwyk.com/ClipKIT/) with the ‘-m kpic’ algorithm (a strategy that retains sites that are either parsimony-informative or constant) to reduce compositional heterogeneity. Gene trees were inferred using IQ-TREE with the mixture protein model ‘-m EX_EHO’ and 1,000 UFBoot2 bootstraps^91^.

Genes used for analyses were filtered based on their properties to mitigate common confounding factors in phylogenomic inference^92^. Previous studies have shown that some gene properties are strongly correlated with phylogenetic signal. For alignments, these properties include the number of parsimony-informative sites^93^, relative composition variability (RCV)^94^, and stationary, reversible and homogeneous (SRH)^95^. Tree-based properties include potentially spurious homologs^96^, and average bootstraps support (ABS) values^50^. We calculated three sequence-based properties (number of parsimony-informative sites, RCV, and SRH) and two tree-based properties (potentially spurious sequences, and ABS) to subsample genes and generate matrices for analyses.

The number of parsimony-informative sites of each locus was calculated using default parameters in PhyKIT^97^ (https://jlsteenwyk.com/PhyKIT/usage/index.html), which in an alignment is associated with strong phylogenetic signal^93^, and kept the loci whose number of parsimony-informative sites exceeded 100. Genes with low RCV values are similarly more suitable for phylogenetic analyses, since they harbour less compositional bias. Therefore, we kept genes with RCV values of less than 0.35 using default parameters in PhyKIT. We excluded the SRH assumptions of each locus with ‘--symtest-only’ strategy, *p*-value 0.05, using IQ-TREE v2.0-rc1^98^. The loci with higher *p*-value (usually 0.01–0.1) of symmetry tests should be removed, which means rejected SRH hypotheses. Potentially spurious sequences, i.e., genes with abnormally long branch lengths, were identified using TreeShrink v1.3.7^99^ with an α threshold of ‘-q 0.05’.

Two matrices were generated for phylogenomic analyses. The USCO matrix (USCO75), named as ‘Matrix1’, with 75% completeness, which represents the lowest ratio of taxa for all partitions, was generated using PhyKIT. Genes with ABS values greater than 70 were selected^43^ to generate a new matrix (USCO75_abs70), named as ‘Matrix2’, while maintaining a good number of loci (approximately 50%).

#### Phylogenetic analyses

To account for common sources of systematic errors in phylogenetic inferences, namely missing data^100,101^, paralogy^102^, the heterogeneous nature of amino acid substitution^103,104^, and incomplete lineage sorting (ILS)^92,105^, we conducted phylogenetic analyses with a multi-species coalescent (MSC) model, as well as concatenation-based analyses using heterogeneous models and partitioned maximum likelihood (ML) analyses. The coalescent-based phylogenies were reconstructed in ASTRAL-III v5.6.1^106^ using the MSC model with default parameters to account for ILS, which uses a set of gene trees to estimate branch supports from quartet frequencies^107^. For concatenation-based analyses, we used IQ-TREE and PhyloBayes MPI v1.8b^108^. For partitioned analyses, the best partitioning scheme and substitution models were selected using the relaxed hierarchical clustering algorithm on ModelFinder^109^ implemented in IQ-TREE using the parameters ‘-rclusterf 10’^110^ and ‘--mset LG’. We also conducted unpartitioned analyses to account for different aspects of heterogeneity in the substitution process. To account for across-site compositional heterogeneity in a ML framework, analyses were conducted with the C60+F+R^111,112^ and PMSF (LG+C60+F+R) models in IQ-TREE that partition the sites of an alignment into 60 compositional categories. For PMSF trees, the corresponding ASTRAL trees with Matrix1 (H1_guide-tree), Matrix1-con (H4_guide-tree), partitioned ML tree with Matrix1 (H2_guide-tree), and C60 tree with Matrix1 (H3_guide-tree) were treated as the initial guide trees. 1,000 SH-aLRT replicates^113^ and 1,000 UFBoot2 bootstraps were calculated for all node supports in the ML analyses. To account for across-site compositional heterogeneity in a Bayesian setting, we combined the unconstrained category (CAT) and general time reversible (GTR) substitution matrices (CAT-GTR) in PhyloBayes. Six independent Markov Chain Monte Carlo (MCMC) of 1,164‒5,317 generations sampled every one generation were run. The two chains converged on a similar topology, except for incongruences within Paraneoptera and Polyneoptera, likely due to narrower taxon sampling for these clades. The phylogenetic relationships within both groups have been the subject of previous studies^114–116^ and do not affect our main results and conclusion, which concern the early-diverging hexapods. We removed the first 3,000 generations as the burn-in. All trees were visualized and edited with iTOL v4^117^. The gCF and sCF were calculated by using IQ-TREE with the option ‘--scf 100’, to quantify genealogical concordance in phylogenomic datasets^118^.

#### Inconsistent genes and gene-wise phylogenetic signal conflict in phylogenomic data matrices

Topological conflict is widespread in phylogenomics^119^. We estimated the gene-wise phylogenetic signal (ΔGLS) for each gene by comparing the sequence alignment to the ML concatenated species (T1: Protura + ((Collembola + Diplura) + Insecta); inferred by C60 model based on Matrix1) and the ASTRAL tree (T2: Collembola + (Protura + (Diplura + Insecta)); inferred by MSC model based on Matrix1). Furthermore, we also calculated gene-wise quartet scores (ΔGQS), which estimates the number of congruent quartets recovered from each gene tree compared to the concatenated species tree. The inconsistent genes in Matrix1 and Matrix2, i.e., those with ΔGLS>0 (a higher log-likelihood score for T1 versus T2) or ΔGQS<0 (a lower quartet score for T1 versus T2), or vice versa, were identified and filtered. Therefore, the two new matrices, USCO75_consistent-genes (referred to as ‘Matrix1-con’) and USCO75_abs70_consistent-genes (referred to as ‘Matrix2-con’), were generated. These two matrices were subjected to the same phylogenomic analyses as those outlined above for Matrix1 and Matrix2.

#### Topology tests

A total of four different hypotheses (H1–4; Fig. 3) were generated with our four analysed matrices. The hypotheses were compared, with all four matrices, using the approximately unbiased (AU), weighted Kishino-Hasegawa (WKH), and weighted Shimodaira-Hasegawa (WSH) tests under the C60+F+R and LG+PMSF(C60) (H3_guide-tree) models in IQ-TREE. The four hypotheses were as follows: H1: Collembola + (Protura + (Diplura + Insecta)); H2: (Collembola + Protura) + (Diplura + Insecta); H3: Protura + ((Collembola + Diplura) + Insecta); H4: (Protura + (Collembola + Diplura)) + Insecta.

#### Cross-validation analyses

We conducted Bayesian cross-validation (CV) in PhyloBayes^108^ to compare the fit of the CAT-GTR and LG models for Matrix2-con. A random subsample of 10,000 sites for ten replicates were run, and each replicate containing 9,000 sites for training the model and 1,000 sites for computing the cross-validation log-likelihood scores. Two independent runs were run for 5,000 generations of each replicate, with parameters and trees sampled every one generation, and the first 2,000 generations were discarded as burn-in. The Wilcoxon test was conducted using R v4.3.1 to compare the difference of cross-validation log-likelihood scores between the two models. Custom script and commands are available from GitHub https://github.com/xtmtd/Phylogenomics/tree/main/basal_hexapods/scripts.

#### Phylogeny without outgroup taxa

To test whether outgroup sampling affected reconstructions of deep nodes in the ingroup, we performed analyses of Matrix2-con with the three outgroup species excluded, using IQ-TREE. First, an unrooted tree was inferred using reversible models^98^. The partition file followed the same results of phylogeny with outgroup included. Second, a rooted tree with linked non-reversible models^98^ was inferred. A rooted tree with linked non-reversible models was inferred to measure the confidence in the root placement. These unrooted and rooted trees were compared with those from phylogenies with the outgroup included.

#### Highlights

- Protura are ‘basal’ to all other hexapods
- Genome-scale analyses show that Diplura and Collembola form a clade
- Previous contentious results likely result from restricted taxon sampling and inadequate substitution modelling

## Notes

### Competing Interest Statement

The authors have declared no competing interest.

